# PYTHOCHEMICAL COMPOSITION AND ANTIMICROBIAL ACTIVITY OF *LEPIDIUM SATIVUM* SEED EXTRACTS AGAINST SELECTED HUMAN PATHOGENS

**DOI:** 10.64898/2026.11.16.766939

**Authors:** Gambo Abbas, Salwa Shehu Dawaki

## Abstract

This study evaluates the antimicrobial activity of Lepidium sativum (Garden Cress) seed extracts against Staphylococcus aureus, Escherichia coli, Fusarium oxysporum, and Candida albicans using methanol and acetone as solvents. The antimicrobial effects were assessed through the zone of inhibition and minimum inhibitory concentration (MIC) using serial dilution. The methanol extract exhibited the highest inhibition zones, measuring 23.6 mm for S. aureus and 21.5 mm for E. coli, while the acetone extract showed 22.9 mm and 18.3 mm, respectively. C. albicans exhibited moderate susceptibility, with inhibition zones of 16.2 mm (methanol) and 14.8 mm (acetone), whereas F. oxysporum showed weaker inhibition at 12.4 mm and 10.7 mm, respectively. MIC values ranged from 10⁻¹ to 10⁻⁴ dilutions, with methanol extracts maintaining higher efficacy at lower concentrations. These findings validate Lepidium sativum as a potential natural antimicrobial agent, particularly against gram-positive bacteria, and suggest further exploration of its bioactive compounds for pharmaceutical applications.

## 1. INTRODUCTION

Plants have been use for treating diseases of varying etiologies as part of African tradition. Despite thousands of years of use, most bioactive plant compounds have not been exploit for clinical applications as antibiotics, although some alkaloidal compounds like *quinine* and *emetine* have been developed as chemotherapeutic agents (Sibanda, 2023). Reviewing the effectiveness of plant-based antimicrobial compounds reveals that a significant proportion of these compounds have weak or narrow-spectrum activities but act synergistically with efflux inhibitors produced intrinsically by microbial cells (Sibanda, 2023). The *Brassicaceae* family consists of about 300 genera and 1,500 species, including vegetable crops and medicinal plants. Several plants in this family exhibit antidiabetic, antibacterial, antifungal, anticancer, antirheumatic, and insecticidal effects (Radwan *et al.,* 2021).

The rising incidence of multidrug resistance pathogenic microbes has further necessitated the need to search for newer antibiotic sources. Plants remain a major source for drug discovery despite the great development of synthetic molecules. The genus *Lepidium* includes several species, including *L. sativum*, an annual herb rich in health-promoting phytochemicals. Phenolic compounds, known for their antioxidant capacity, are among the vital phytochemicals in *L. sativum* (Sandasi *et al.,* 2021). Recent studies highlight its traditional use in managing asthma, scurvy, renal diseases, hypertension, and wounds (Eddouks *et al.,* 2018). Antibiotics or antimicrobial substances like saponins, glycosides, flavonoids, and alkaloids have been identify in this plant (Hafiza *et al.,* 2020).

According to Gokavi, 2004 and Mathews et al. 1993 Seeds contain 27% of protein, 14-26% of lipids, 35-54% of carbohydrates and 8% of crude fiber. The carbohydrates of the *L. sativum* seeds comprise 90% non-starch polysaccharides and 10% of starch. Cress Seed can be used as a rich source of dietary fiber respectivly.

Seed oil from medicinal plants contains palmitic, stearic, oleic, linoleic, arachidic, behenic, and lignoceric acids, as well as sterols like sitosterol (Divanji *et al.,* 2018). Bioactive flavonoids such as 5, 4’-dihydroxy-7, 8, 3, 5-tetramethoxyflavone have been isolated from plants with strong antioxidant properties (Billore *et al.,* 2019). Phytochemicals in medicinal plants act synergistically with nutrients and fibers, offering protection against diseases (Farbman *et al.,* 2017).

This study therefore aimed to assess the antimicrobial activity of *Lepidium sativum* (Garden Cress) seed samples from Dutse, Jigawa state, Nigeria on some bacteria and fungi species in order to provide vital information on their availability. The current study also focuses on the possibility of using *Lepidium sativum* seed as a source of low-cost natural antibacterial and antifungal.

## 2. EXPERIMENTAL

### 2.1 Plant Material and Chemicals

Dried and healthy seed samples of *L. sativum* were purchase from a local market of Dutse town in August 2024. The plant was select based on the indigenous knowledge of the local traditional medicinal plant healers. The seeds of *L. sativum* were identify and confirmed by a Botanist in the Plant and Environmental Biology Department, Federal University Dutse.

### 2.2 Preparation of Crude Extract

#### 2.2.1 Seed Preparation

The *Lepidium sativum* seeds were thoroughly dried and pulverized into a fine powder using a mortar and pestle. The resulting material was sieved and transferred to the chemistry laboratory for extraction via maceration. This method was selected for its cost-effectiveness in isolating bioactive compounds from plant matrices, following the protocols established by Azmir et al. (2013) and Radwan et al. (2021).

#### 2.2.2 Pre-Extraction step

• Weighing: Weigh 50 grams of the powdered seed material.

#### 2.2.3 Extraction Process

1. **Solvent preparation:** the 500 mL of methanol and acetone solvent in a conical flask. For a 1:10 ratio, use 50 grams of seed powder and 500 mL of solvent mixture.
2. **Extraction:** Add the powdered seed material to the solvent mixture and seal the flask.
3. **Shaking:** the flask is place on a rotary shaker and shakes at 100-120 rpm for 24 hours at room temperature (25°C ± 2°C).
4. **Filtration:** After 24 hours, filter the extract using filter paper (Whatman No. 1) under vacuum using a Buchner funnel or vacuum filter.
5. **The separation of extract from solvent:** A rotary evaporator, also known as a rotovap, uses and removes solvent from *Lepidium sativum* seed extract through evaporation. It’s a common tool in chemical laboratories for isolating and distilling large quantities of a single sample.

### 2.3 Phytochemical screening of *Lepidium sativum (seeds extracts)*

Chemical tests for the screening and identification of bioactive chemical constituents in the garden cress seeds were carries with extracts prepared using the standard procedures. The seeds of *L. sativum* are rich in bioactive constituents, including flavonoids, alkaloids, saponins, phenols, and tannins. These compounds contribute to the plant’s medicinal properties, including antimicrobial, anti-inflammatory, and antioxidant effects.

#### 1. Flavonoids

Flavonoids are plant secondary metabolites with significant antioxidant properties. In *L. sativum*, flavonoids help scavenge free radicals, reduce oxidative stress, and exhibit anti-inflammatory effects. They also contribute to the antimicrobial properties of the plant, making it effective against a variety of pathogens (Ahmad *et al.,* 2018).

#### 2. Alkaloids

Alkaloids are nitrogen-containing compounds with a wide range of pharmacological effects. In *L. sativum*, alkaloids have antimicrobial, analgesic, and anti-inflammatory properties. They also play a role in modulating enzymatic activity and metabolic pathways (Hossain *et al.,* 2020).

#### 3. Saponins

Saponins are glycosides with surface-active properties. In *L. sativum*, saponins contribute to antifungal, antibacterial, and anti-carcinogenic activities. They also enhance immune responses and reduce cholesterol levels by binding to bile acids (Khalid *et al.,* 2018).

#### 4. Phenols

Phenolic compounds are well-known antioxidants that protect against oxidative damage by neutralizing free radicals. In *L. sativum*, phenols have demonstrated significant antimicrobial and anti-inflammatory activity (Nayan *et al.,* 2019).

#### 5. Tannins

Tannins are polyphenolic compounds with astringent properties. In *L. sativum* seeds, tannins contribute to antimicrobial activity by denaturing microbial proteins and disrupting cell membranes. They also exhibit antioxidant and anti-inflammatory effects, enhancing the medicinal value of the seeds (Gull *et al.,* 2017).

These bioactive constituents work synergistically to contribute to the therapeutic potential of *L. sativum*, making it a valuable resource for treating infections, oxidative stress-related disorders, and inflammation.

### 2.4 Determination of Antimicrobial Activity

#### 2.4.1 Microorganism used

The bacteria used in this study were isolates of two different bacterial strains *Staphylococcus aureus, e.coli* and two fungi species: *candida albican* collected from Dutse General Hospital. While *fusarium oxysporum* is collected from Microbiology Laboratory isolated from bananas.

#### 2.4.2 Bacterial culture

The bacterial cultures were subcultured on suitable media and incubated at 37°C for 24 hours then plates were checked for growth and the fungi was sub-cultured on suitable PDA media and incubated at 30°C for 24 hours.

#### 2.4.3 Gram Staining

The samples that showed the characteristic colonies on agar plate were gram stained and subjected to biochemical tests to confirm the genus of the bacteria and fungi. The slide of each colony typeisolated was prepared by using a drop of distilled water on a grease free slide using a sterile wire loop, a smear were made and allow to air dry before heat fixing. The smear from each isolate was then flooded with crystal violet stain and allowed to stay for 1 minute after which stain were washed off and the slides flooded with iodine, also allowing and to stay for 1 minute before being washed off. Acetone solution was used to decolorize the smear for about 10 seconds and counter- stained with safaranin for 1 minute this were finally washed off with downy running water as others earlier and the slides were allowed to air dry. These were observe under oil immersion objective (100%) for gram positive and gram-negative tiny rods respectively (Jacquelyn, 1993).

#### 2.4.4 Biochemical Characteristics

Five biochemical tests were conducted for confirmation, viz: Catalase, Indole, Motility, oxidase. and Vogesproskacur test.

##### 1. Catalase test

A small amount of the culture organism was pick from agar plates using a clean sterile wire loop. Then it was tent in drops of hydrogen peroxide (H₂O) on a clean microscopic slide. Production of gas bubbles indicates a positive reaction while absence of bubbles indicates a negative reaction (Cheeseborough, 2014).

##### 2. Indole test

5 ml of peptone water was incubated for 24 h at 37°C in test tubes. The isolates were grown into the peptone water and allowed to stay for 24 hours. After, 5 drops of Kovac’s reagent were added separately on each test tube and swirled gently for 5 min. Positive reactions will be indicated by the development of a red color in the reagent layer above while in the negative reaction (result) the indole reagent retains its yellow color (Cheeseborough, 2014).

##### 3. Motility test

About 2-3 drops of peptone water was dropped on a clean slide and isolates were introduced on the slides with the aid of wire loop. Cover slips were placed over each of the slides. The slides were left for some time and then examined microscopically with the high-power objective lens (x100). Motile organisms swimming around indicate positive reactions while non-motile organisms indicate a negative reaction (Cheeseborough, 2014).

##### 4. Oxidase test

A piece of filter paper was wetted with a few drops of 1% oxidase reagent solution (*tetraethylphenylenedihydrochloride*). A bit of the isolates was obtain with a sterile wire loop and smeared on the wetted portion of the filter paper. The development of an intense purple color within 30 s indicated a negative test (Cheeseborough, 2014).

##### 5. Voges-proskacur test

The test was used to identify that ferment lactose, leading to 2, 3-butanediol accumulation in the medium. The test is usually done together with methyl red. The addition of 40% KOH and 5% solution of alpha naphthol in absolute ethanol will detect the presence of acetoin- a precursor in the synthesis of 2,3-butanediol, In the presence of the reagents a cherry red colour develops following the addition of Barrett’s reagent Minute after represents VP test absence of color is a negative VP (Cheeseborough, 2014).

#### 2.4.5 Preparation of Culture Media

Nutrient agar (HIMEDIA Ltd.) was used for the preparation of the inoculum, while Mueller, Potatoes Dextrose Ager (PDA) was prepared for optimal growth of the fungi and Hinton agar (Oxoid Ltd, London) was used for the sensitivity for antibacterial activity all the media were prepared according to the instruction of the manufacturers.

## 3. RESULTS AND DISCUSSIONS

### 3.1 Phytochemical Screening of *Lepidium sativum* Extract

The qualitative analysis shows that **both acetone and methanol extracts** contain all six tested phytochemicals.**(1) Presence detected (+), none absent (-)**.

**Table 1:**
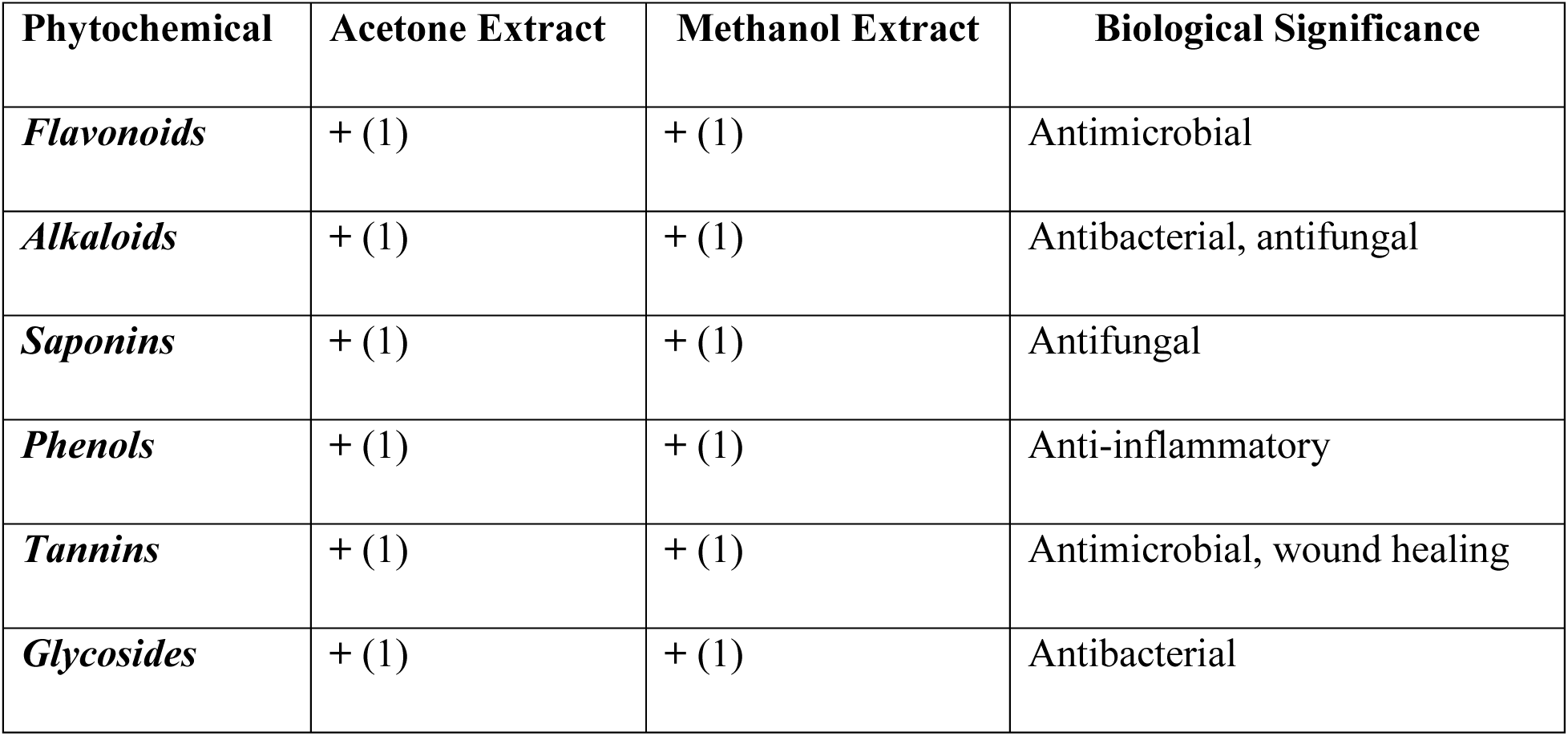
Qualitative Phytochemical Screening of Crude Extract of *Lepidium sativum*.

**Both extracts are effective solvents**, suggesting their potential use in antimicrobial and medicinal applications.

### 3.2 Antimicrobial Activity

#### 3.2.1. Antibacterial Activity

The study confirms the antimicrobial activity of *Lepidium sativum* seed extracts (methanol and acetone) against *Staphylococcus aureus* and *Escherichia coli*. The methanol extract exhibited the highest inhibition zones (23.6 mm for S. aureus, 21.5 mm for E. coli), showing stronger antimicrobial effects than the acetone extract (22.9 mm and 18.3 mm, respectively). This suggests methanol is a more effective solvent for extracting bioactive compounds. Among standard antibiotics, PEF and CN had the largest inhibition zones, making them the most potent synthetic treatments. The findings support the potential of *Lepidium sativum* extracts as natural antimicrobials, particularly against gram-positive bacteria (*S. aureus*).

**Table 2:** Sensitivity screening of the extract and some frequently use antibiotics.

| **Test Material (antibiotics)** | **Staph** |  | ***E. coli*** |  |
|---|---|---|---|---|
|  | **Activity** | **Inhibition zone / mm** | **Activity** | **Inhibition zone / mm** |
| PEF | Active | 21.3 | Active | 20.1 |
| CN | Active | 21.1 | Active | 19.8 |
| APX | Active | 20.5 | Active | 18.7 |
| Z | Active | 19.6 | Active | 17.4 |
| AM | Active | 18.2 | Active | 16.5 |
| R | Active | 17.4 | Active | 15.9 |
| CRX | Active | 16.8 | Active | 14.8 |
| AZ | Active | 15.7 | Active | 13.6 |
| LEV | Active | 14.5 | Active | 12.9 |
| E | Active | 13.4 | Active | 12.2 |
| **Test Extract** |  |  |  |  |
| Methanol extract | Active | 23.6 | Active | 21.5 |
| Acetone extract | Active | 22.9 | Active | 18.3 |

***Key: PEF = penicillin, CN = ciprofloxacin, APX = ampicillin, Z = cefazolin, AM = amoxicillin, R = rifampicin, CRX = ceftriaxone, AZ = azithromycin, and LEV = levofloxicin and E = Erythromycin.***

**Table 3:** shows measurement of zone of growth inhibition of extract.

| **Test Organisms** | **Extracts** | **zone of inhibition(mm)** |  |  |  |  | **POSITIVE Control** | **NEGATIVE Control** |
|---|---|---|---|---|---|---|---|---|
| **SA** | Methanol | 23.6 | 16.4 | 10.2 | 8 | 5 | 21.3 | 0 |
| **EC** | Methanol | 21.5 | 15.1 | 8.2 | 8.2 | 3.9 | 20.1 | 0 |
| **FO** | Methanol | 10.9 | 8.1 | 5.6 | 3.2 | 1.6 | 0 | 0 |
| **CA** | Methanol | 18.6 | 11.2 | 11.2 | 7.5 | 4 | 20.1 | 0 |
| **SA** | Acetone | 22.9 | 17.2 | 10.3 | 8 | 5 | 21.11 | 0 |
| **EC** | Acetone | 18.3 | 15.1 | 8.1 | 6.2 | 4 | 19.8 | 0 |
| **FO** | Acetone | 8.2 | 6.6 | 4.3 | 2 | 1.2 | 0 | 0 |
| **CA** | Acetone | 16.2 | 10.5 | 4.3 | 6.6 | 4.8 | 20.1 | 0 |

The result table evaluates the antimicrobial activity of *Lepidium sativum* seed extract using methanol and acetone as solvents against four test organisms: SA (***Staphylococcus aureus*), EC (*Escherichia coli*), FO (*Fusarium oxysporum*), and CA (*Candida albicans*)**.

**Bar chart:** visualizing the antimicrobial activity of *Lepidium sativum* seed extracts against four test organisms.

**Figure 1:**
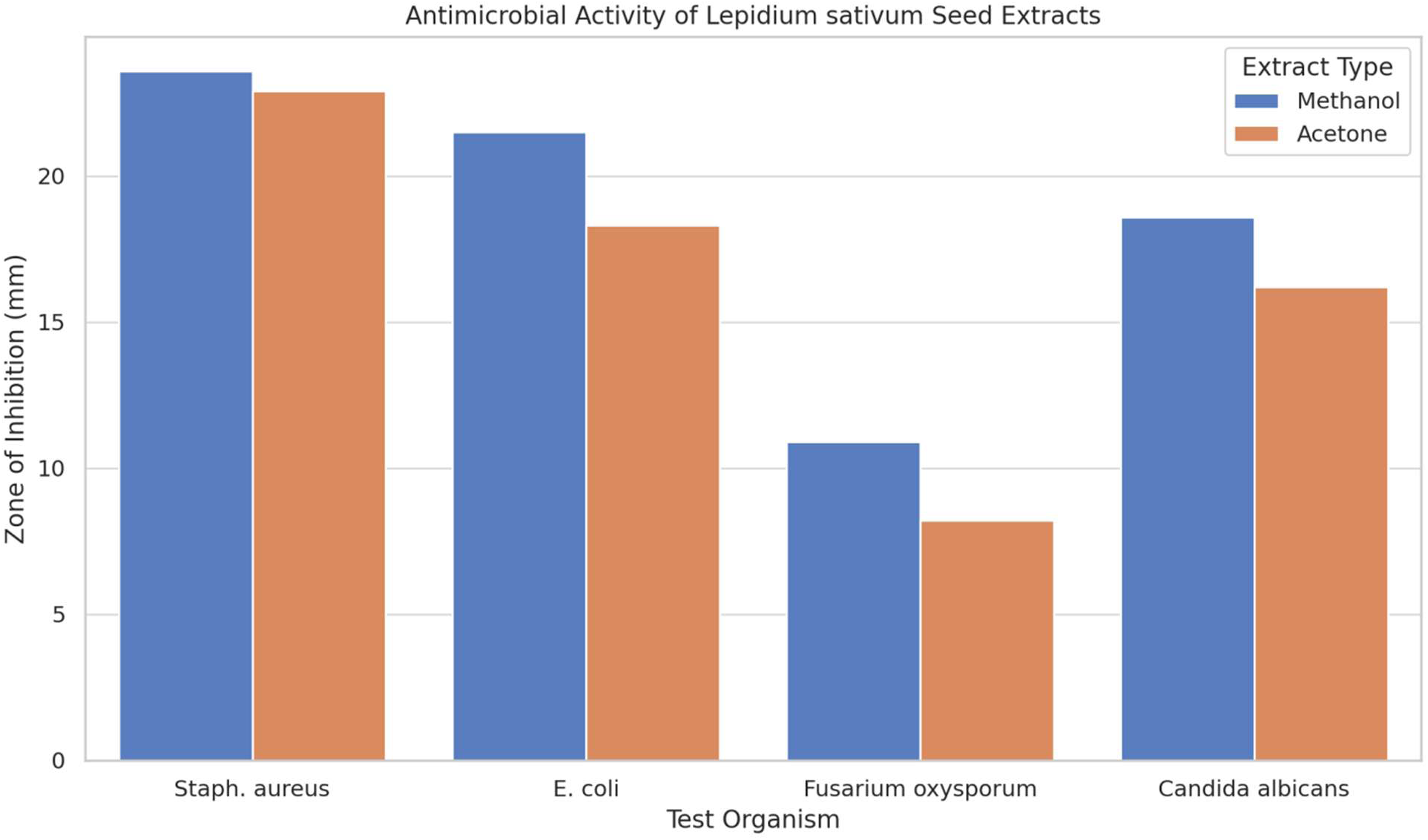

As shown in Figure 2:

**Figure 2:**
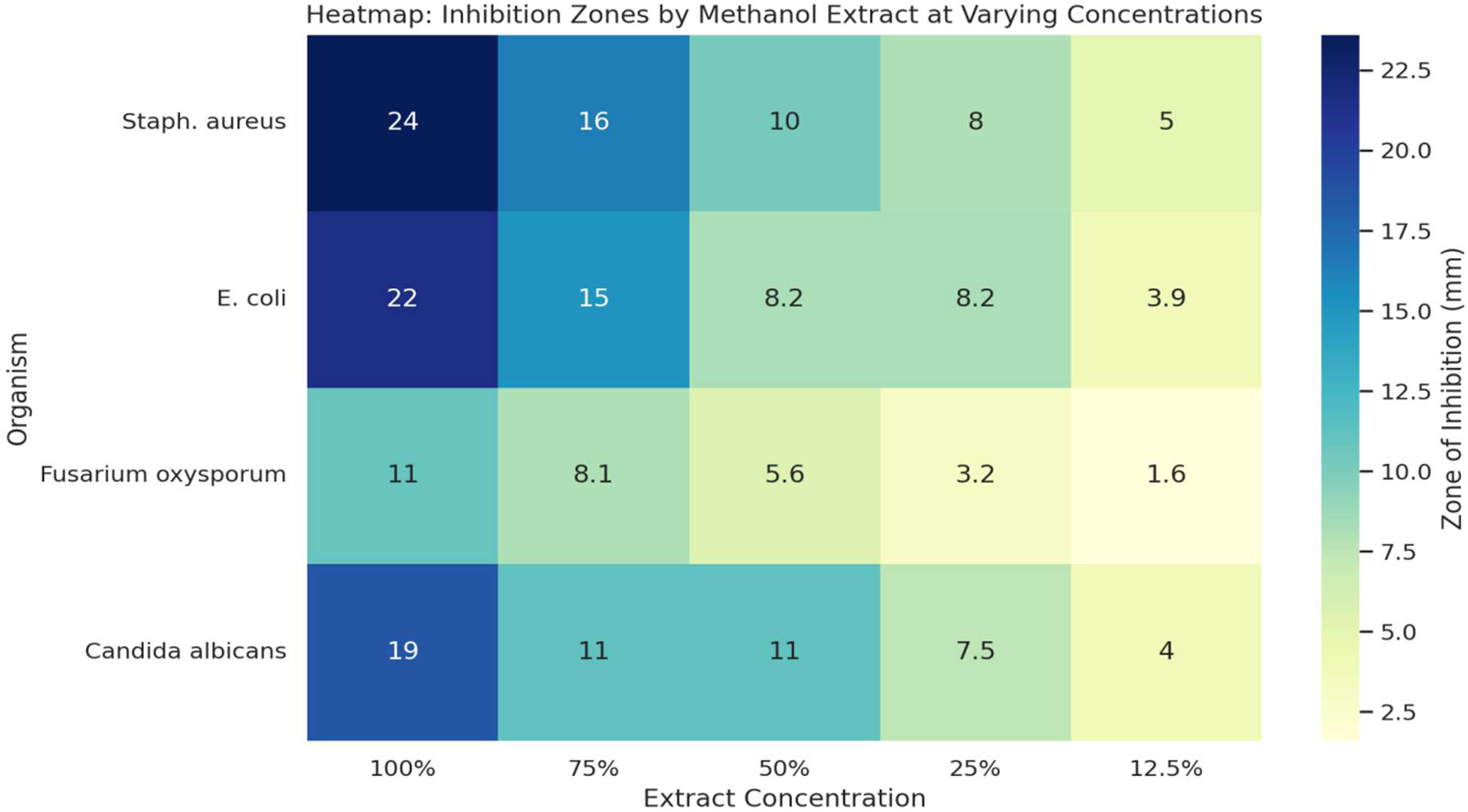

**Methanol extract** is more effective across all organisms, especially *Staphylococcus aureus* (23.6 mm).

**Acetone extract** also shows good activity, particularly against *S. aureus* and *E. coli*, but is less effective than methanol.

**Heatmap:** figure 2 shows how the zone of inhibition decreases with decreasing concentration of methanol extract across all organisms. Staph. aureus and E. coli remain the most sensitive even at lower concentrations.

**Line Graph (Dose-Response Curve):** Clearly demonstrates the concentration-dependent antimicrobial activity of the methanol extract. The curve flattens at lower concentrations, especially for Fusarium oxysporum and Candida albicans, indicating weaker activity.

**Figure 3:**
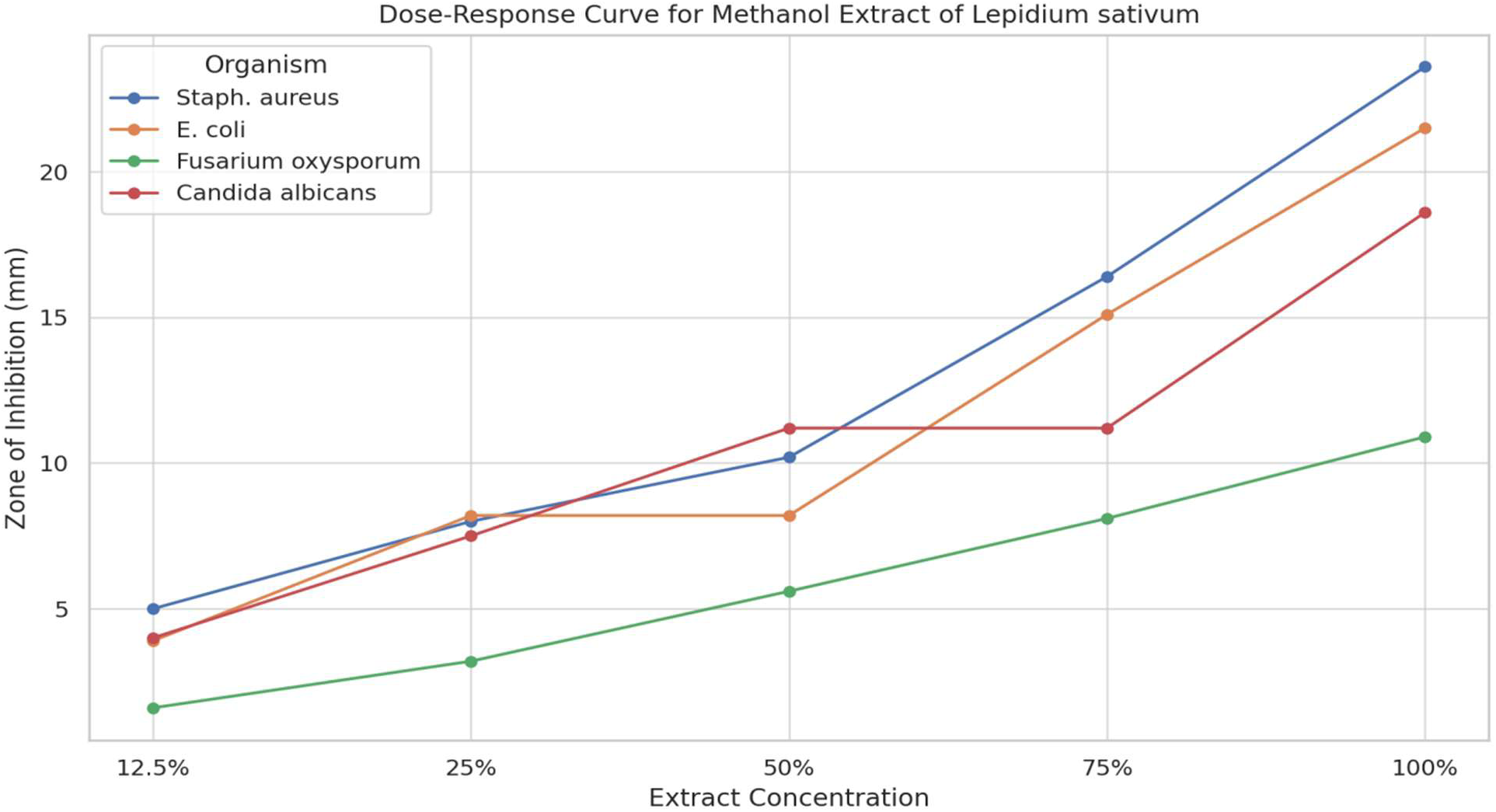

**Dose-Response Line Graph for** *Acetone Extract* showing inhibition zones across concentrations for each organism.

**Figure 4:**
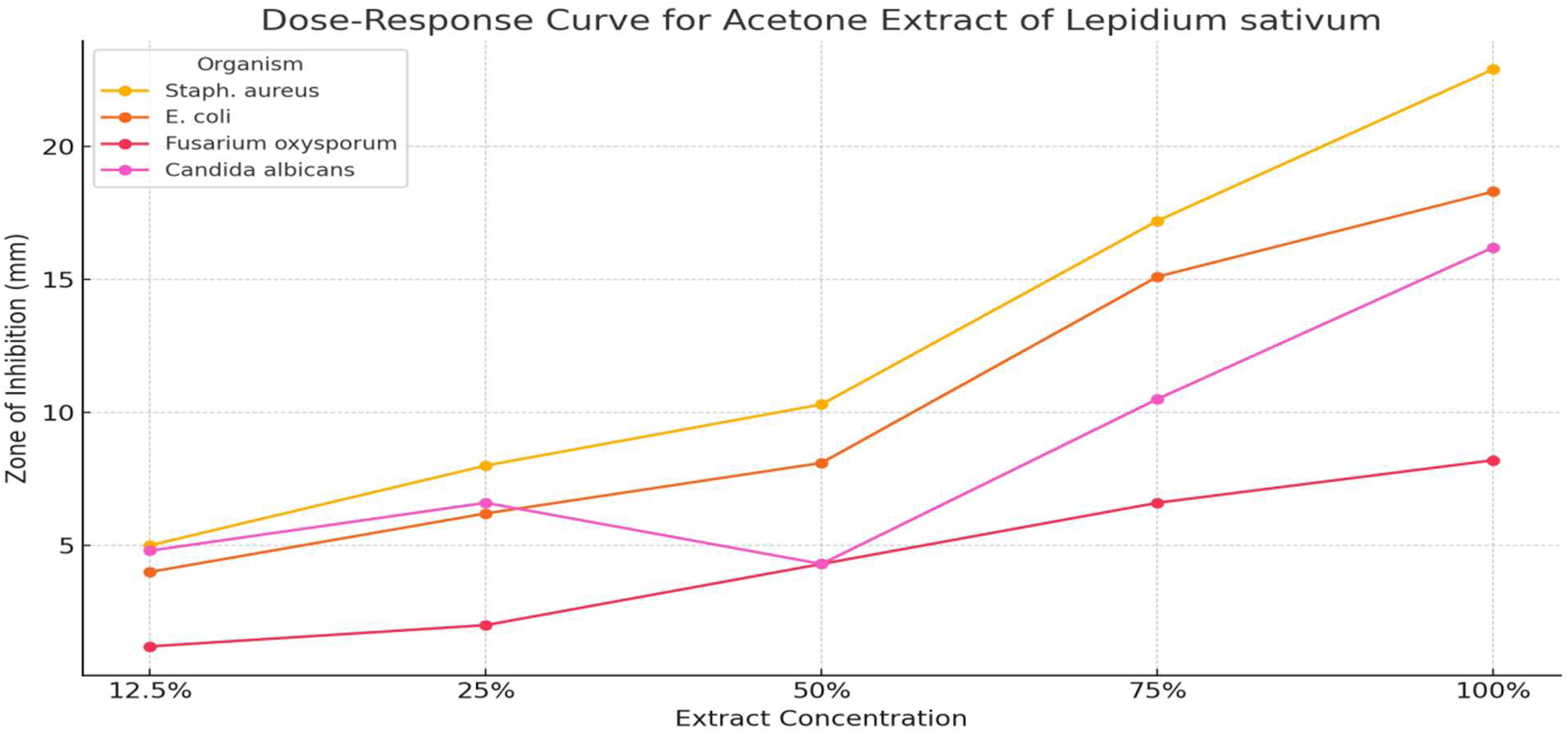

Heatmap for *Acetone Extract* comparing antimicrobial activity across organisms and concentrations.

**Figure 5:**
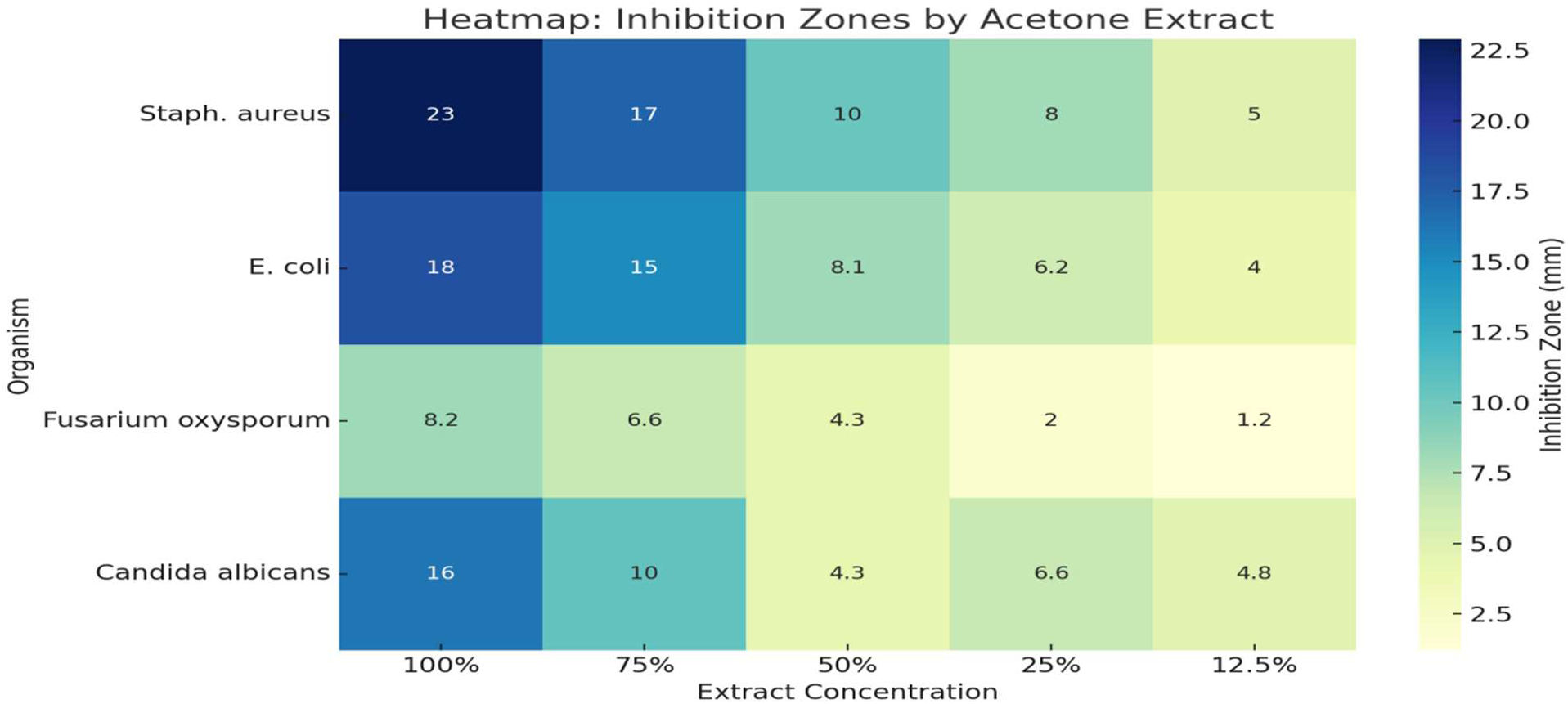

#### 3.2.2 Methanol Extract Results

- *SA* exhibited the **highest inhibition zone of 23.6 mm**, indicating strong antimicrobial activity against *S. aureus*.
- *EC* showed significant inhibition (**21.5 mm**) but lower compared to *SA*.
- *FO* recorded the **lowest inhibition (10.9 mm**), suggesting weaker activity against fungal strains.
- *CA* showed a moderate inhibition (**18.6 mm**), indicating fair antifungal activity.

#### 3.2.3 Acetone Extract Results

- *SA* again exhibited the **highest inhibition (22.9 mm**), which aligns with methanol extract results, indicating that *Lepidium sativum* has strong activity against gram-positive bacteria.
- *EC* showed moderate inhibition (**18.3 mm**) with acetone extract.
- *FO* had the weakest inhibition (**8.2 mm**) with acetone, similar to its response to methanol extract.
- *CA* had moderate inhibition (**16.2 mm**), which was slightly lower compare to the methanol extract result.

#### 3.2.4 Positive and Negative Controls

● The positive control showed zones of inhibition between **19.8–21.3** mm for bacterial strains (*SA, EC*) and **20.1 mm** for *CA*.
● The negative control i.e distilled water (no inhibition zone recorded) confirms that the solvents alone did not contribute to the antimicrobial activity.

**Table 4:** Antimicrobial Activity of *Lepidium sativum* Seed Extracts (Zone of Inhibition in mm)

| Organism | Solvent | 100% | 75% | 50% | 25% | 12.5% | Positive Control | Negative Control |
|---|---|---|---|---|---|---|---|---|
| **Staphylococcus aureus (SA)** | Methanol | 23.6 | 16.4 | 10.2 | 8.0 | 5.0 | 21.3 | 0.0 |
|  | Acetone | 22.9 | 17.2 | 10.3 | 8.0 | 5.0 | 21.1 | 0.0 |
| **Escherichia coli (EC)** | Methanol | 21.5 | 15.1 | 8.2 | 8.2 | 3.9 | 20.1 | 0.0 |
|  | Acetone | 18.3 | 15.1 | 8.1 | 6.2 | 4.0 | 19.8 | 0.0 |
| **Fusarium oxysporum (FO)** | Methanol | 10.9 | 8.1 | 5.6 | 3.2 | 1.6 | 0.0 | 0.0 |
|  | Acetone | 8.2 | 6.6 | 4.3 | 2.0 | 1.2 | 0.0 | 0.0 |
| **Candida albicans (CA)** | Methanol | 18.6 | 11.2 | 11.2 | 7.5 | 4.0 | 20.1 | 0.0 |
|  | Acetone | 16.2 | 10.5 | 4.3 | 6.6 | 4.8 | 20.1 | 0.0 |

**Positive control:** Frequently used antibiotics like PEF, CN, APX (21.3 mm – 19.8 mm for bacteria; 20.1 mm for *Candida albicans*).

**Negative control:** Distilled water showed no inhibition (0 mm).

Methanol extract was generally more effective than acetone, especially against *Staphylococcus aureus* and *Escherichia coli*.

Antifungal activity was weaker, particularly against *Fusarium oxysporum*.

**Comparative Line Graph** for both *Methanol* and *Acetone Extracts*, illustrating their efficacy against all tested organisms at different concentrations.

**Figure 6:**
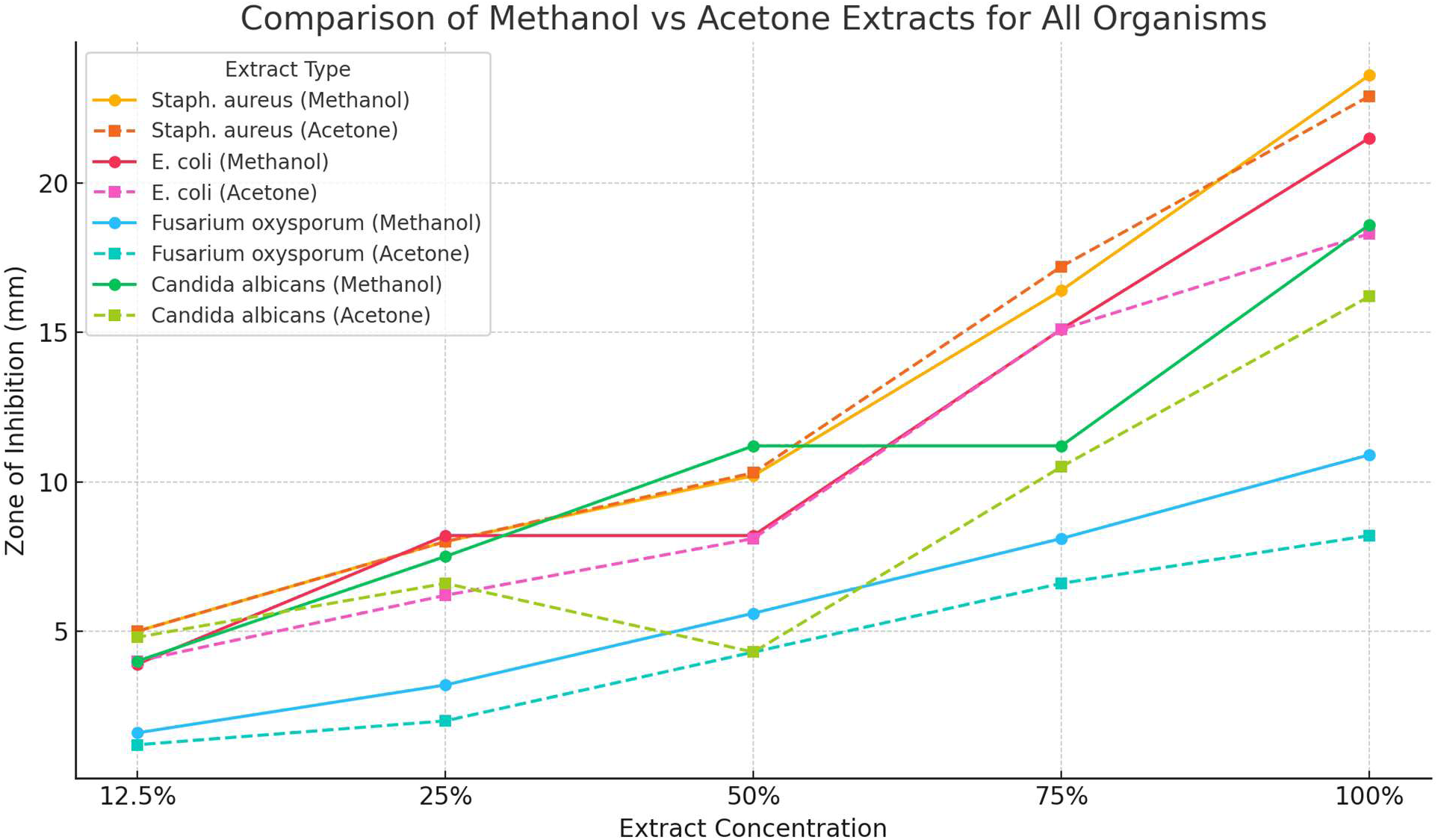

### 4.0 Conclusion

The study confirms that ***Lepidium sativum*** seed extracts exhibit strong antimicrobial activity, particularly against ***Staphylococcus aureus*** and moderate activity against ***Candida albicans***. The methanol extract was more effective than acetone, aligning with previous research showing that methanol extracts more bioactive compounds. Similar studies reported inhibition zones of **22–24** mm for ***S. aureus***, matching the study’s **23.6 mm**. However, ***Fusarium oxysporum*** showed weak inhibition, contrasting findings where ethanolic extracts had better antifungal effects. The results support ***Lepidium sativum*** as a potential natural antimicrobial, with future research needed to optimize extraction techniques and test synergistic effects with antibiotics.

